# Quantifying feedback among traits in coevolutionary models

**DOI:** 10.1101/2025.02.28.640831

**Authors:** Xiaoyan Long, Jussi Lehtonen, Jonathan Henshaw

## Abstract

Phenotypic traits rarely evolve in isolation. Instead, multiple traits typically interact, resulting in complex coevolutionary dynamics. Such dynamics can be predicted using mathematical frameworks such as adaptive dynamics and quantitative genetics. Selection gradients play a crucial role in these frameworks, describing the direction and strength of selection and predicting evolutionary trajectories and potential endpoints. Current theory focuses mainly on analysing how traits change in response to selection. However, selection gradients also change over time as traits evolve. For a complete understanding of coevolutionary dynamics, it is consequently essential to examine how trait changes feed back to influence the selection environment. Here, we develop a general framework for investigating coevolutionary feedback between traits and selection gradients. Rather than relying on verbal interpretation of causal relationships, our approach explicitly quantifies the various pathways by which traits and selection gradients influence one another. Our framework can be applied both to adaptive-dynamic models and to quantitative-genetic models under the weak selection limit. We illustrate our approach with three examples that showcase its potential to deepen our understanding of established models.

## Introduction

In nature, it is rare for one trait to evolve independently of the rest of an organism’s phenotype. Instead, multiple traits typically interact to determine an organism’s fitness, resulting in often complex coevolutionary dynamics (Svensson et al., 2021; Bank, 2022). Here are three examples of this ubiquitous phenomenon, taken from our own field of behavioural ecology: First, sex role divergence often involves the coevolution of male and female traits, including parental care, mate choice and mating competition (Andersson, 1994; Fromhage & Jennions, 2016; Henshaw et al., 2019; Henshaw et al., 2022; Long & Weissing, 2023). Second, natal dispersal is suggested to coevolve with prosocial behaviours (e.g., helping other members in a group; see Hochberg et al., 2008; Powers et al., 2011; Mullon et al., 2018; McNamara & Leimar, 2020). Across species, a high dispersal rate is often associated with a low level of prosociality (for potential evolutionary explanations, see Hochberg et al., 2008 and Mullon et al., 2018). Third, in group-foraging species foraging strategies coevolve with vigilance against predators (Ranta et al., 1998; Giraldeau & Caraco, 2000). In particular, it has been suggested that producers, who primarily focus on searching for food, may face higher costs when scanning for predators, whereas scroungers, who rely on producers for food resources, can allocate more time and energy to vigilance (Mathot & Giraldeau, 2008; Harten et al., 2018).

Our understanding of coevolutionary dynamics depends critically on theoretical models based on a range of approaches, including population genetics (Kirkpatrick, 1982; Nagylaki, 1992; Revathi Venkateswaran et al., 2021), quantitative genetics (Lande, 1981; Roff, 1997; Mead & Arnold, 2004), adaptive dynamics (Fawcett et al., 2011; Henshaw et al., 2019; Dashtbali et al., 2024) and individual-based simulations (Baldauf et al., 2014; Long & Weissing, 2023; Waffender & Henshaw, 2023; Long et al., 2024). Here, our primary focus is on the adaptive-dynamic framework. By separating the processes of mutation, selection and demographic or ecological change, adaptive dynamics enables us to focus on phenotypic, rather than genetic, evolution. This facilitates models of complex trait interactions, including frequency-dependent selection and demographic structure (Pen & Weissing, 2001; Kuijper et al. 2012). In adaptive-dynamic models, rare mutations of small effect are assumed to arise against the background of a monomorphic resident population (where all other individuals share the same phenotype). The relationship between mutational effects and mutant fitness is quantified as a selection gradient, which describes the direction and strength of selection acting on a trait. Selection gradients can then be used to predict evolutionary trajectories and the likely end points of evolution (e.g., evolutionarily stable strategies, see Geritz et al., 1998, McGill & Brown, 2007, Dercole & Rinaldi, 2008).

Selection gradients are a key determinant of evolutionary change in adaptive dynamics and many other modelling frameworks (e.g., Lande & Arnold, 1983; Taylor, 1996). Crucially, selection gradients themselves also change over evolutionary time, for two reasons. First, selection is often frequency-dependent, so that the relationship between trait values and fitness (i.e., the fitness function) changes along with the population trait distribution. Second, since selection gradients depend on the statistical relationship between trait values and fitness within a population, they depend on the current trait distribution even when the fitness function is held fixed. Selection’s dependence on the trait distribution in turn has important implications for coevolution, because it induces a feedback loop: Selection causes traits to change, leading to changes in the current trait distribution, which can subsequently alter the strength and direction of selection. Further, just as selection on one trait can indirectly cause changes in other traits due to genetic correlations (Lande & Arnold, 1983), changes in the distribution of one trait can affect selection on other coevolving traits, because fitness is often determined by an interaction between traits (Svensson et al., 2021; Bank, 2022). Our current mathematical toolkit mainly focusses on analysing the first part of this feedback loop (i.e., the effects of selection on trait evolution). For a complete understanding of coevolutionary dynamics, however, it is essential to also understand how changes in traits feed back to influence the selection environment.

Here, we develop a general framework for analysing feedback between traits and selection gradients in coevolutionary models. Our goal is to enhance our understanding of trait coevolution by providing a formal tool to examine how traits interact and subsequently shift the selection environment. Instead of relying solely on verbal explanations of coevolutionary dynamics, our approach offers a means of quantifying feedbacks. Our framework can be applied to adaptive-dynamic models and also to quantitative genetic models under the assumption of weak selection. We illustrate these applications via three examples, revealing our framework’s potential to enrich our understanding of even well-established models.

## General approach

### Adaptive dynamics for a single trait

We begin by outlining the fundamental assumptions of adaptive-dynamic models of evolutionary change, as encapsulated by the ‘canonical equation’ (Dieckmann & Law, 1996; Champagnat et al., 2001). In adaptive dynamics, the resident population is assumed to be genetically monomorphic at any given point in time. Mutations arise rarely and cause only small deviations from the resident phenotype. If a mutant trait value outperforms the resident strategy in fitness terms, there is a non-zero probability that it invades the resident population, and eventually completely replaces the resident strategy. Since mutations are rare, it is assumed that each invasion is completed before a new mutation arises. This process typically continues until the population reaches an equilibrium state, at which point no further mutant invasions are possible. The dynamics of this process are captured by the canonical equation of adaptive dynamics (Dieckmann & Law, 1996). In a one-trait model with a resident trait *z*, the canonical equation takes the form:

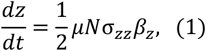

where *μ* is the mutation rate of the trait at birth; *N* is the size of the resident population; σ_*zz*_ is the variance in the phenotypic effects of mutations (given by 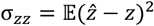, where 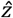 is the trait value of a mutant); the selection gradient 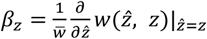 describes the strength and direction of selection on the trait *z* (where 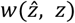 denotes the fitness of a mutant with trait 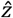 in a monomorphic resident population with trait *z*, and 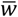 indicates the mean fitness of resident individuals); and the factor 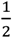 reflects that, under directional selection, only half of the mutations are beneficial and thus can contribute to evolutionary change. The factor 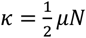 indicates the total number of mutations that can contribute to evolutionary change per generation. Thus, the one-trait canonical equation can also be written as

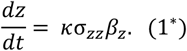

### Feedback analysis with one evolving trait

We will principally be interested in feedback acting on two or more coevolving traits. However, we will first introduce some of the necessary concepts in the setting of a single evolving trait, where things are much simpler. As shown in Figure 1, the selection gradient *β*_*z*_ determines the direction and scales the speed of evolutionary changes on the trait *z* (black arrow) and, reciprocally, changes in the trait value can feed back to influence the selection gradient (grey arrow). Using the chain rule of calculus, we can break down the rate of change in the selection gradient as follows:

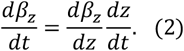

**Figure 1.**
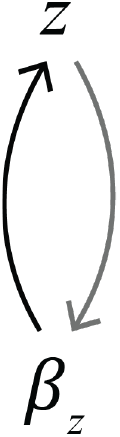
Feedback loop between a single trait *z* and its selection gradient *β*_*z*_ in an adaptive-dynamic model. The black arrow indicates changes in the trait value due to selection, whereas the grey arrow represents changes in the selection gradient due to changing trait values.

Plugging equation (1^*^) into (2) then yields:

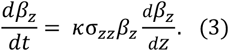

Equation (3) describes the feedback loop acting on the selection gradient (Fig. 1) in quantitative terms. To see qualitatively how such feedback affects selection, we need to examine the combination of the selection gradient *β*_*z*_ and its feedback term 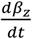 (Fig. 2). A positive selection gradient indicates selection for trait values that are larger than the current resident strategy. When feedback is also positive, it intensifies this selection, making the selection for a larger trait even stronger. We refer to this as ‘reinforcing feedback’. On the other hand, if the selection gradient is positive, but feedback is negative, it weakens selection for larger trait values, which we term ‘inhibiting feedback’. Analogous considerations apply when the selection gradient is negative: Negative feedback on a negative selection gradient reinforces selection, whereas positive feedback inhibits it. Thus, feedback can either amplify or dampen selection on a trait, depending on whether it aligns with or opposes the direction of selection gradient (see Examples below).

**Figure 2.**
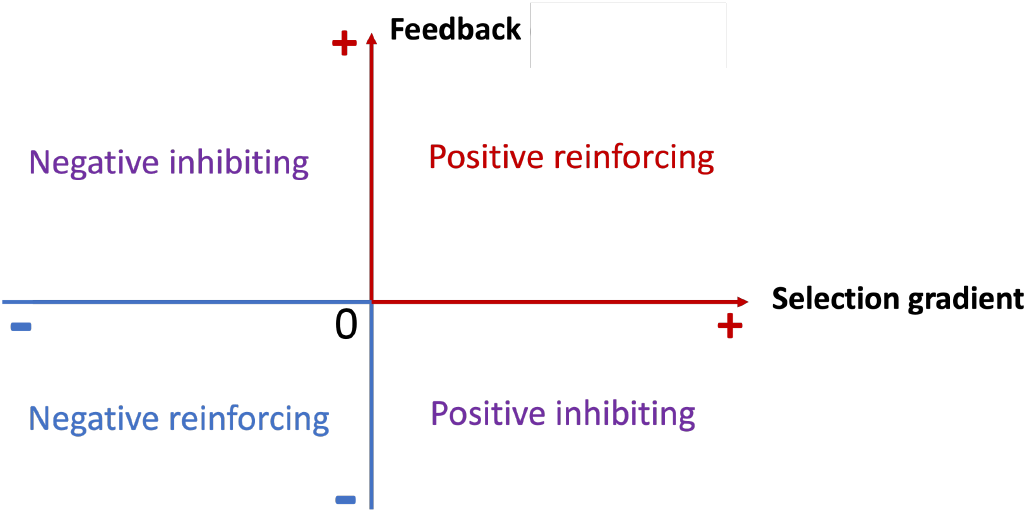
How does feedback influence the strength of selection on a trait? In this coordinate plot, the x-axis represents the selection gradient, while the y-axis represents the feedback (or feedback components) acting on the selection gradient (e.g., equation (3)). When both the selection gradient and feedback are positive (top-right quadrant), selection favors mutants with trait values larger than the current resident value, and feedback acts to increase the intensity of this selection (positive reinforcing selection). When the selection gradient is positive but feedback is negative (bottom-right quadrant), selection again favors larger trait values, but feedback reduces the strength of this selection (positive inhibiting selection). In the bottom-left quadrant, where both the selection gradient and feedback are negative, selection favors smaller trait values and feedback increases the strength of selection (negative reinforcing selection). Finally, in the top-left quadrant, when the selection gradient is negative but feedback is positive, selection again favors smaller trait values, but feedback acts to weaken selection (negative inhibiting selection).

### Adaptive dynamics of two coevolving traits

When multiple traits coevolve in a single species, the evolutionary dynamics become more complicated, because traits can interact with one another. Here we start with a model with two traits, *x* and *y*. In this case, the canonical equation takes the form:

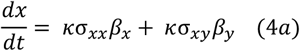

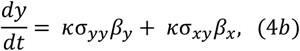

which can be written more compactly as

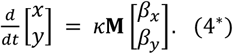

Here, 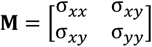 is the variance-covariance matrix of the phenotypic effects of new mutations, and 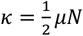 is defined as above. Importantly, σ_*xy*_ represents the mutational covariance between the two traits (i.e., the covariance between the effects of a random mutation on *x* and *y*). Such mutational covariance can affect the direction of evolution. If *σ*_*xy*_ ≠ 0, then selection acts on *x* both directly (quantified by *β*_*x*_) and indirectly via selection on *y* (*β*_*y*_) and the mutational correlation between the two traits (*σ*_*xy*_).

### Feedback analysis with two coevolving traits

For ease of understanding, we start with the assumption that there is no mutational correlation between the coevolving traits (i.e., the off-diagonal elements of the matrix **M** equal zero), meaning that each trait is only under direct selection. This assumption is widely made in applications of adaptive dynamics. As a result, equation (4*a*, 4*b*) simplifies to:

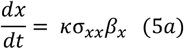

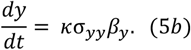

We now develop tools for studying coevolutionary feedback between two coevolving traits. In the absence of mutational correlation, such feedback arises via the following two-step process (Fig. 3A): (1) Traits change due to direct selection (black arrows in Fig. 3A). (2) The selection environment (i.e., the strength and direction of selection) changes due to the shifts in trait values (grey arrows in Fig. 3A). In the two-trait model, each selection gradient, *β*_*x*_ and *β*_*y*_, can be understood as a function of the current resident trait values, *x* and *y*. Similar to equation (2), we can break down the selection gradients *β*_*x*_ and *β*_*y*_ using the chain rule, as follows:

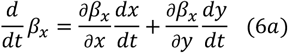

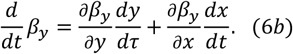

**Figure 3.**
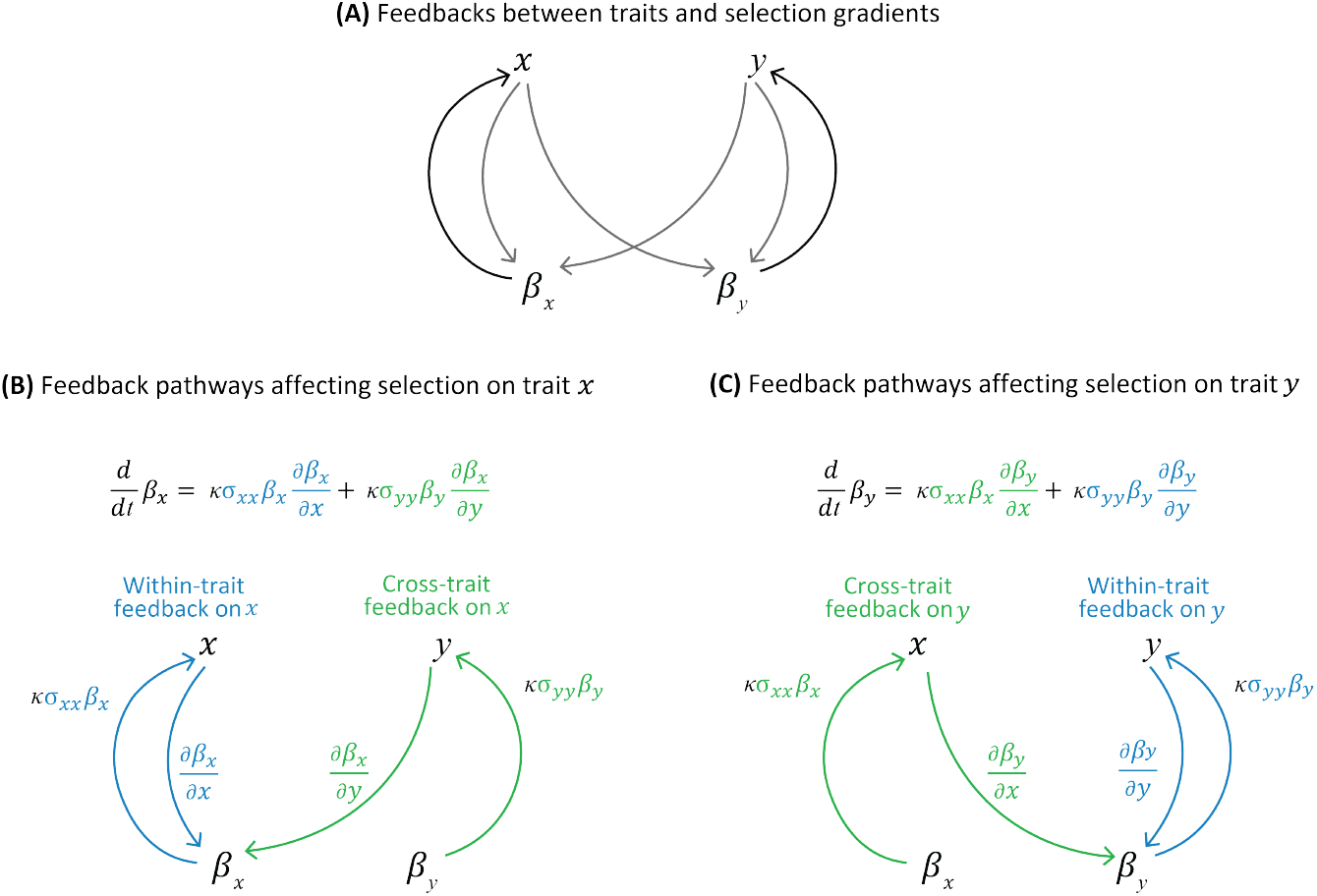
Feedback analysis for two coevolving traits in the absence of mutational covariance. **(A)** Feedback pathways among the four key elements – the traits *x* and *y* and the selection gradients *β*_*x*_ and *β*_*y*_ – when there is no correlation in mutational effects between the two traits. In this case, the selection gradients *β*_*x*_ and *β*_*y*_ influence the respective traits *x* and *y* (black arrows). Changes in the values of each trait feed back to influence both selection gradients (grey arrows). **(B)** Feedback analysis of the selection gradient *β*_*x*_ in the absence of mutational covariance. Feedback on *β*_*x*_ consists of two components: First, direct selection on the focal trait *x* feeds back to influence selection *β*_*x*_ on the same trait (blue arrows). This feedback is referred as ‘within-trait feedback’ and quantified as 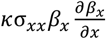. Only this type of feedback is present in a one-trait model (cf. Fig. 1). Second, direct selection on the other trait *y* affects *β*_*x*_ (green arrows). This feedback is referred as ‘cross-trait feedback’ and quantified as 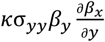. **(C)** Feedback analysis of the selection gradient *β*_*y*_ in the absence of mutational covariance. Again, feedback on *β*_*y*_ consists of two components. First, direct selection on the focal trait *y* feeds back to influence selection *β*_*y*_ on the same trait (blue arrows, ‘within-trait feedback’: quantified as 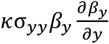). Second, direct selection on the other trait *x* affects *β*_*y*_ (green arrows, ‘cross-trait feedback’: quantified as 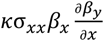).

Plugging equations (5*a*) and (5*b*) into (6*a*) and (6*b*) then yields:

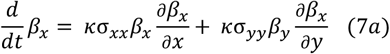

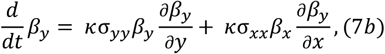

which can be written more compactly as:

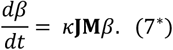

where 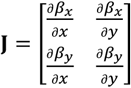 is a Jacobian matrix, 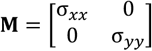 and 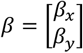.

The Jacobian matrix **J** plays an important role in the existing theory on multivariate adaptive dynamics, in particular in relation to the convergence stability of an evolutionarily stable strategy (ESS) (i.e., whether nearby resident strategies will converge to the ESS over evolutionary time, see Geritz et al., 1998; McNamara & Leimar, 2020). In general, an ESS is convergence stable if the eigenvalues of the matrix product **JM**, evaluated at the ESS trait values, have negative real parts. Since the matrix **M** is typically unknown, an additional criterion is often useful: Convergence stability is guaranteed for all fixed variance-covariance matrices **M** if the matrix **J** + **J**^*T*^ is negative definite at the ESS (McNamara & Leimar, 2020).

Equations (7*a*) and (7*b*) describe how changes in the traits *x* and *y* affect the selection gradients *β*_*x*_ and *β*_*y*_ in the absence of mutational covariance, respectively. To understand how changes in each trait feed back to alter the selection gradient *β*_*x*_ acting on the trait *x*, we can analyze the structure of equation (7*a*) (see Fig. 3B). In fact, there are two feedback components: (1) Direct selection on the focal trait *x* can affect selection *β*_*x*_ acting on the same trait (‘within-trait feedback’, 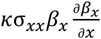: blue arrows in Fig. 3B). This is the only type of feedback in the one-trait model presented above (cf. Fig. 1) (2) Changes to the other trait *y* may likewise affect the selection gradient *β*_*x*_ (‘cross-trait feedback’, 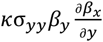: green arrows in Fig. 3B). Within-trait feedback describes how direct selection on the focal trait affects its selection gradient, whereas cross-trait feedback indicates how interactions between traits influence selection on the focal trait. The structure of feedback on the selection gradient *β*_*y*_ (equation 7*b*) is analyzed analogously in Figure 3C.

To gain a better understanding of how trait change influences selection, we can analyze the combination of the selection gradient and each feedback component for a given trait (see Fig. 2). Similar to the one-trait model, each feedback component can either strengthen or weaken the selection on a focal trait. The overall strength and magnitude of feedback are determined by the sum of the feedback components. We can consequently compare the magnitude of each feedback component to determine which component has a more pronounced effect on the strength of selection acting on the focal trait (see Examples 2 and 3).

The feedback analysis becomes more complicated when mutations have correlated effects on multiple traits (Fig. 4). In this case, a trait may change because it is under direct selection, or because it is genetically correlated with other traits that are under selection, as shown in equations (4*a*) and (4*b*).

**Figure 4.**
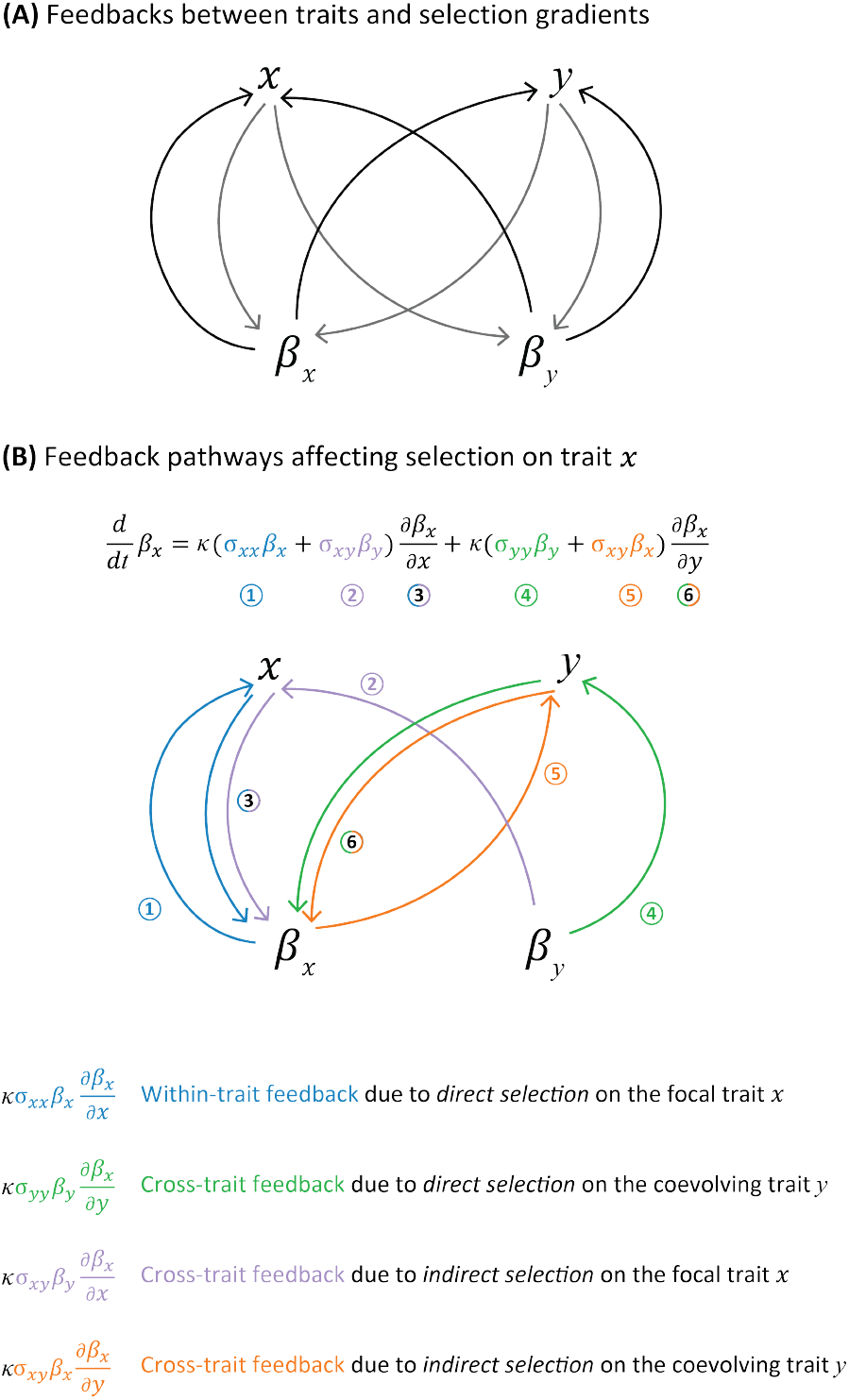
Feedback analysis with mutational covariance between traits. **(A)** Feedback pathways between trait values and selection gradients in the presence of mutational covariance. In this case, both selection gradients *β*_*x*_ and *β*_*y*_ influence each trait via direct and indirect selection (black arrows). Changes in the values of each trait feed back to influence both selection gradients (grey arrows). **(B)** Feedback analysis of the selection gradient *β*_*x*_ in the presence of mutational correlation. Feedback on the selection gradient consists of four components. First, direct selection on the focal trait *y* feeds back to influence selection *β*_*x*_ on the same trait (blue arrows; quantified as 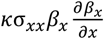). Second, direct selection on the other trait *x* affects *β*_*x*_ (green arrows; quantified as 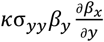). Only these two types of feedback exist in the absence of mutational covariance (cf. Fig. 3). Third, indirect selection on the focal trait *x* due to its correlation with the trait *y* influences *β*_*x*_ (purple arrows; quantified as 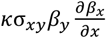). Fourth, indirect selection on the trait *y* due to its correlation with the focal trait *x* can also influence *β*_*x*_ (orange arrows; quantified as 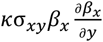).

Plugging these equations into (6*a*) and (6*b*) then yields:

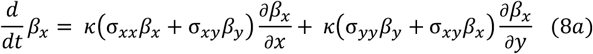

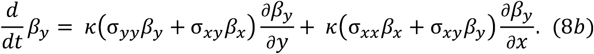

which can again be written more compactly as equation (7^*^), this time with a fully general mutational variance-covariance matrix 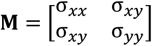.

Equations (8*a*) and (8*b*) describe how changes in the traits *x* and *y* affect the selection gradients *β*_*x*_ and *β*_*y*_, respectively, in the presence of mutational covariance. Here we take equation (8*a*) as an example to understand how various feedback pathways affect selection on *x* (Fig. 4B). When there is mutational correlation between the two traits, there are four feedback components: (1) Direct selection on the focal trait *x* can affect selection on the same trait (‘within-trait feedback’, 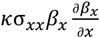: blue arrows in Fig. 4B). (2) The focal trait *x* may evolve due to selection on *y*, in combination with the mutational correlation between *x* and *y*, which in turn affects the selection gradient *β*_*x*_ (‘cross-trait feedback due to indirect selection on the focal trait’, 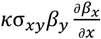: purple arrows in Fig. 4B). Changes in the other trait *y* may likewise affect the selection gradient *β*_*x*_. Such changes may result either from (3) direct selection on *y* (‘cross-trait feedback due to direct selection on the coevolving trait’, 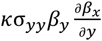: green arrows in Fig. 4B), or from (4) selection on *x* in combination with a mutational correlation between *x* and *y* (‘cross-trait feedback due to indirect selection on the coevolving trait’, 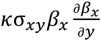: orange arrows in Fig. 4B).

Here, the ‘within-trait feedback’– the first of the four feedback components – describes how direct selection on the focal trait affect its selection gradient (the same as the blue arrows in Fig. 3B and 3C), whereas all the other three feedback components indicate how interactions between the traits influence selection on the focal trait.

### Feedback analysis of multivariate evolution

We focused above on the case of two coevolving traits. The extension to multiple traits (*z*_1_, *z*_2_, …, *z*_*n*_) can be made by setting 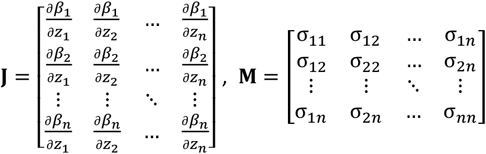 and 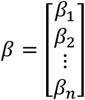 in equation (7^*^). Our arguments are thus general and apply also to coevolution among more than two traits. As in the two-trait case, within-trait feedback (e.g., blue arrows in Figs. 3 and 4) represents how direct selection on the focal trait affects its selection gradient, whereas all other feedback components describe how interactions between traits influence selection on the focal trait.

### Feedback analysis in discrete-time adaptive dynamics

In adaptive dynamics, the continuous-time evolutionary process can also be approximated by a discrete-time model. Such approximations are often useful for simulating evolutionary trajectories in practice (e.g., detailed steps are provided in the Appendix of Lehtonen & Kokko, 2011; see also Fromhage & Jennions, 2016; Henshaw et al., 2019). In a one-trait model with a resident trait *z*, the trait value after a discrete time step Δ*t* is approximated by the recursion equation

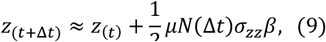

For simplicity, we write 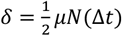 for the total number of mutations that contribute to evolutionary change over the time step Δ*t*. The discrete-time approximation of the canonical equation can then be written as

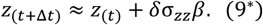

Similarly, in a two-trait model with resident traits *x* and *y*, the discrete-time approximation of the canonical equation for each trait is given by

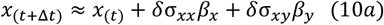

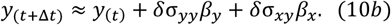

with 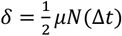 defined as above.

The feedback analysis in the discrete-time model is very similar to the continuous-time model shown above. In the discrete-time model, we can approximate the fitness gradients (*β*_*x*_’, *β*_*y*_’) at time *t* + Δ*t* using a first-order Taylor polynomial, as evolution is assumed to proceed gradually. Writing (*x*′, *y*′) for the trait value at time *t* + Δ*t*, we have:

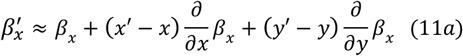

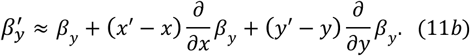

Plugging in equations (10*a*) and (10*b*) then yields:

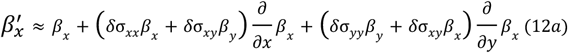

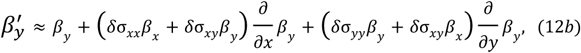

We can write the equation of (12) in a more compact way:

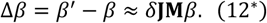

with Jacobian matrix 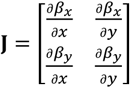, variance-covariance matrix 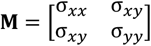, and selection gradients 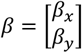. This equation can be analyzed analogously to equation (7^*^) above.

### Feedback analysis in the weak selection limit of quantitative genetics

The above approach to selection feedback analysis – which is based on discrete approximations to adaptive-dynamic models – can be readily adapted to a class of quantitative-genetic models that assume weak selection. In the weak selection limit, each trait is assumed to distribute sharply around the mean value, so that the fitness function is nearly linear over the range of the trait value (Iwasa et al., 1991). Under this assumption, the equations for evolutionary dynamics for quantitative genetic models are analogous in form to the discretized canonical equation of adaptive dynamics (Iwasa et al., 1991; Leimar, 2009). Therefore, we can also analyze the coevolutionary feedback between multiple traits in quantitative genetics under weak selection. To apply our approach, trait variation at any given time point should be sufficiently small (explicitly, it should be small relative to the radius of curvature of the fitness function). Typically, this assumption is taken to mean that traits are distributed sharply around the mean. However, the assumption can also hold when traits deviate substantially from the mean, as long as the fitness function remains approximately linear over the range of trait variation.

To see how this approach works, we start with a quantitative genetic model of a single trait *z*. The change in mean trait value 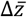 from the current generation to the next generation is given by

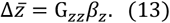

where G_*zz*_ is the genetic variance of the trait *z*, and *β*_*z*_ is the selection gradient, defined in this context as 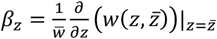, where 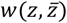 is the fitness of an individual with trait value *z* in a population with mean trait value 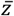, and 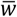 indicates the mean fitness of the population. Note that under the weak selection assumption, the definition of the selection gradient coincides with that of adaptive dynamics (cf. Taylor, 1996; Dieckmann & Law, 1996).

Similarly, the coevolution of two traits, *x* and *y*, can be described as

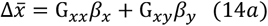

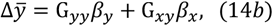

or more compactly as

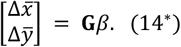

where 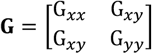 is the additive genetic variance-covariance matrix. The **G**-matrix and the **M**-matrix share similar roles in describing variance and covariance in traits. The **M**-matrix describes mutational effects, where the diagonal elements indicate variances in the mutational effects on traits, and the off-diagonal elements show covariances in mutational effects due to pleiotropy (Jones et al., 2007; Arnold et al., 2008). The **G**-matrix, on the other hand, reflects standing genetic variation, with its diagonal consisting of additive genetic variances for traits and its off-diagonal elements capturing additive genetic covariances arising from pleiotropy and linkage disequilibrium (Pigliucci, 2006; Arnold et al., 2008). Despite these similarities, **M**- and **G**-matrices operate on different time scales. The **M**-matrix operates on shorter timescales, highlighting the effects of new mutations as they occur. In contrast, the **G**-matrix is more dynamic and reflects evolutionary processes operating over short, medium, and long-term timescales, incorporating the cumulative effects of mutation, selection, and genetic drift over many generations (Arnold et al., 2008).

Similar to the discrete-time model in adaptive dynamics, Taylor series can be used to approximate the fitness gradients (*β*_*x*_’, *β*_*y*_’) at the next time step given the trait value (*x*′, *y*′), analogously to equations (11–12). This gives us:

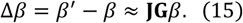

with Jacobian matrix 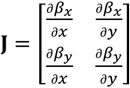, genetic variance-covariance matrix 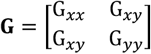, and selection gradient 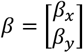. Feedback on selection can then be analyzed analogously to above.

## Examples

To illustrate the application of our feedback analysis, we present three examples. Beginning with an adaptive-dynamic framework, we will explore scenarios involving a single trait (Example 1) and two coevolving traits (Example 2). Lastly, we will present an example that utilizes the framework of quantitative genetics (Example 3). With these examples, we aim to showcase the versatility and efficacy of our feedback analysis across various evolutionary contexts.

### Example 1: The evolution of offspring size

We first investigate the evolution of a single trait: offspring size (e.g., the size of eggs at laying or offspring at birth in species with no further parental care), building on the framework established by Smith and Fretwell (1974). The original model was formulated as an optimality model, but we consider it here in an adaptive-dynamic framework. We assume that each female has a fixed amount of resource *R* for producing offspring. An individual producing offspring of size *m* can generate 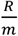 offspring in total. Therefore, smaller offspring can be produced in greater number. On the other hand, an offspring’s survival probability *S* increases with its size according to 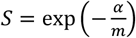, where the parameter *α* scales the relationship between offspring size and survival. There is thus a trade-off between offspring number and survival in this model. The fitness (i.e., the number of surviving offspring) of a rare mutant with offspring size 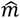 in a resident population with offspring size *m* is given by 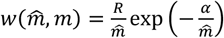. Note that selection is frequency-independent in this simple model, so that 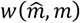 does not depend on the resident strategy *m*. The selection gradient 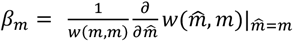 the evolutionary trajectories 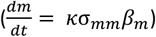 and the feedback acting on the selection gradient 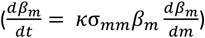 can be derived easily. In this simple model, the feedback is simply the second derivative of the fitness function, which is frequency-independent. Consequently, offspring size evolves towards its optimum at *m* = *α* (Smith and Fretwell 1974). The model’s simplicity allows for a clear understanding of the dynamics driving selection on offspring size over time.

Figure 5 illustrates an exemplary evolutionary trajectory (Fig. 5A), along with the selection gradient (Fig. 5B), and the feedback acting on the selection gradient (Fig. 5C). Since the initial value of *m* exceeds the optimal size in this example, it decreases until reaching the optimal size of *m* = *α* = 1 (Fig. 5A). Consequently, the selection gradient is consistently negative (Fig. 5B). According to Figure 5C, feedback is initially negative (to the left of the dashed vertical line) and then switches to positive (to the right of the dashed vertical line). As offspring size approaches its optimum, both the selection gradient (see Fig. 5B) and feedback term approach zero. By examining the combination of the selection gradient *β*_*m*_ (Fig. 5B) and its feedback term 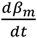 (Fig. 5C), we can understand how feedback affects selection according to Figure 2. At the beginning, both the selection gradient and its corresponding feedback term are negative. Therefore, as offspring size decreases, selection for smaller size becomes stronger (‘negative reinforcing selection’, denoted by the blue background in Fig. 5C), accelerating the evolutionary process towards the optimal offspring size. This self-reinforcing process continues until around generation 7500, from which point the feedback term switches to positive, while the selection gradient remains negative. This reflects ‘negative inhibiting selection’, whereby the selection for smaller offspring weakens as offspring size approaches the equilibrium (purple background in Fig. 5C).

**Figure 5.**
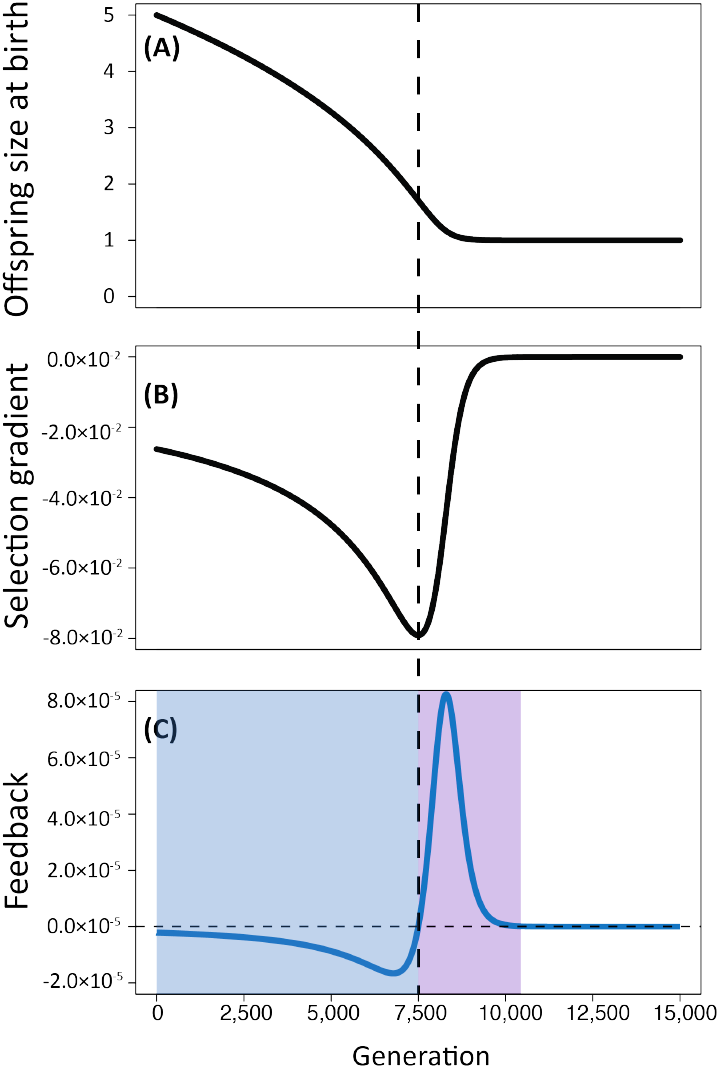
Application of feedback analysis to the evolution of a single trait, offspring size at birth *m*. **(A)** Evolutionary trajectory following the canonical equation (*dm*/*dt* = *κ*σ_*mm*_*β*_*m*_), **(B)** selection gradient *β*_*m*_ and **(C)** feedback 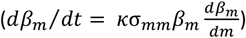 over evolutionary time. The vertical dashed line indicates the time point at which the feedback term switches from negative to positive. In (C), the blue and purple backgrounds indicate the ‘negative reinforcing’ and ‘negative inhibiting’ feedback phases, respectively. Parameter settings: *R* = 1, *α* = 1, *κ* = 0.01 and σ_*mm*_ = 1. Offspring size *m* was initialized at 5. We approximated the evolution trajectory, selection gradient, and feedback numerically using Wolfram Mathematica (see Mathematica code in Supplementary Materials).

### Example 2: The evolution of anisogamy

We next demonstrate the application of our feedback analysis to a model with two coevolving traits: the anisogamy model of Lehtonen and Kokko (2011). In their study, Lehtonen and Kokko explored the impact of gamete competition and gamete limitation on the evolution of anisogamy. The model considers a population consisting of two different mating types, denoted as *x* and *y*, which produce gametes of sizes *m*_*x*_ and *m*_*y*_, respectively. While both gamete types are initially of similar size, disassortative fusion is necessary to produce zygotes (i.e., type *x* can only fuse with type *y* and vice versa). In each local mating group, there are a fixed number of adults, *A*_*x*_ and *A*_*y*_, of each mating type. Each parent in the group possesses a fixed amount of resources *M* for reproduction, resulting in the production of a total number of 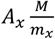 and 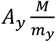 gametes with sizes *m*_*x*_ and *m*_*y*_, respectively. An increase in the values of *A*_*x*_ and *A*_*y*_ corresponds to greater gamete competition. The gametes of each mating type undergo depletion through dying at rates of *μ*(*m*_*x*_) and *μ*(*m*_*y*_), as well as through fertilization occurring at a rate of *γN*_*x*_*N*_*y*_, where *N*_*x*_ and *N*_*y*_ represent the numbers of gametes with sizes *m*_*x*_ and *m*_*y*_ available for fertilization. Here, the parameter *γ* determines the encounter rate between the two types of gametes and is used to modify the gamete limitation in the model. As illustrated in the Supplementary Materials, the dynamics of *N*_*x*_ and *N*_*y*_ are influenced by gamete production, mortality, and fertilization (see equation (*S*1)). Setting equation (*S*1) to zero allows the determination of the equilibrium number of available gametes for each mating type.

Similar to the one-trait model shown above, the fitness of each mating type is defined as the total number of surviving zygotes. Therefore, the fitness of a single mutant which produces gametes of size 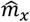 in a resident population with gametes of sizes *m*_*x*_ and *m*_*y*_ is given by 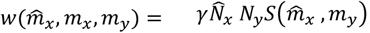, where 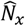 and *N*_*y*_ are the numbers of gametes with sizes 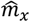 and *m*_*y*_ available for fertilization at equilibrium (see equation (*S*2) in the Supplementary Materials) and 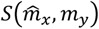 represents the survival probability of the zygotes, which increases with zygote size (given by 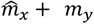). The invasion fitness of a single mutant producing gametes with size 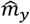 can be derived similarly. Based on the invasion fitness functions, the selection gradients for each gamete type are given by 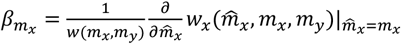 and 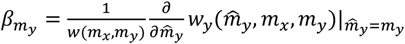 (see Supplementary Materials or Lehtonen and Kokko 2011 for more details).

Lehtonen and Kokko (2011) do not allow for mutational covariance between *m*_*x*_ and *m*_*y*_. Therefore, both traits are only under direct selection. In line with this, the evolutionary trajectories can be described by our equation (5). Feedback acting on each selection gradient (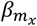 and 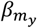) due to changes in gamete sizes can be derived using our equation (7).

Figure 6 shows an exemplary evolutionary trajectory, in which an initially small asymmetry in gamete sizes is magnified, ultimately resulting in the evolution of anisogamy. During this process, each selection gradient is shaped by both within- and cross-trait feedback (see Fig. 3). Gametes of size *m*_*x*_ are initially slightly smaller than opposite-type gametes, gradually decreasing in size until they reach equilibrium (Fig. 6A_1_). Throughout this process, the selection gradient 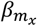 remains negative until it stabilizes at zero (Fig. 6A_2_). Within-trait feedback acting on 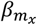 (i.e., changes in 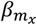 due to direct selection on *m*_*x*_) is initially negative (left of the orange dashed vertical line in Fig. 6A_3_), then switches to positive (right of the orange dashed vertical line in Fig. 6A_3_). Given that the selection gradient is uniformly negative (Fig. 6A_2_), this corresponds to an initial phase of reinforcing within-trait feedback, which acts to intensify the negative selection on *m*_*x*_ (blue background in Fig. 6A_3_), followed by inhibiting within-trait feedback that diminishes selection for smaller *m*_*x*_ (purple background in Fig. 6A_3_). This pattern is analogous to feedback on offspring size in the previous example, which starts as a self-reinforcing process but transitions into a self-inhibiting one as the equilibrium is approached. However, this does not fully describe the course of evolution for *m*_*x*_ in this case, as changes in *m*_*y*_ also impact the selection environment for *m*_*x*_.

**Figure 6.**
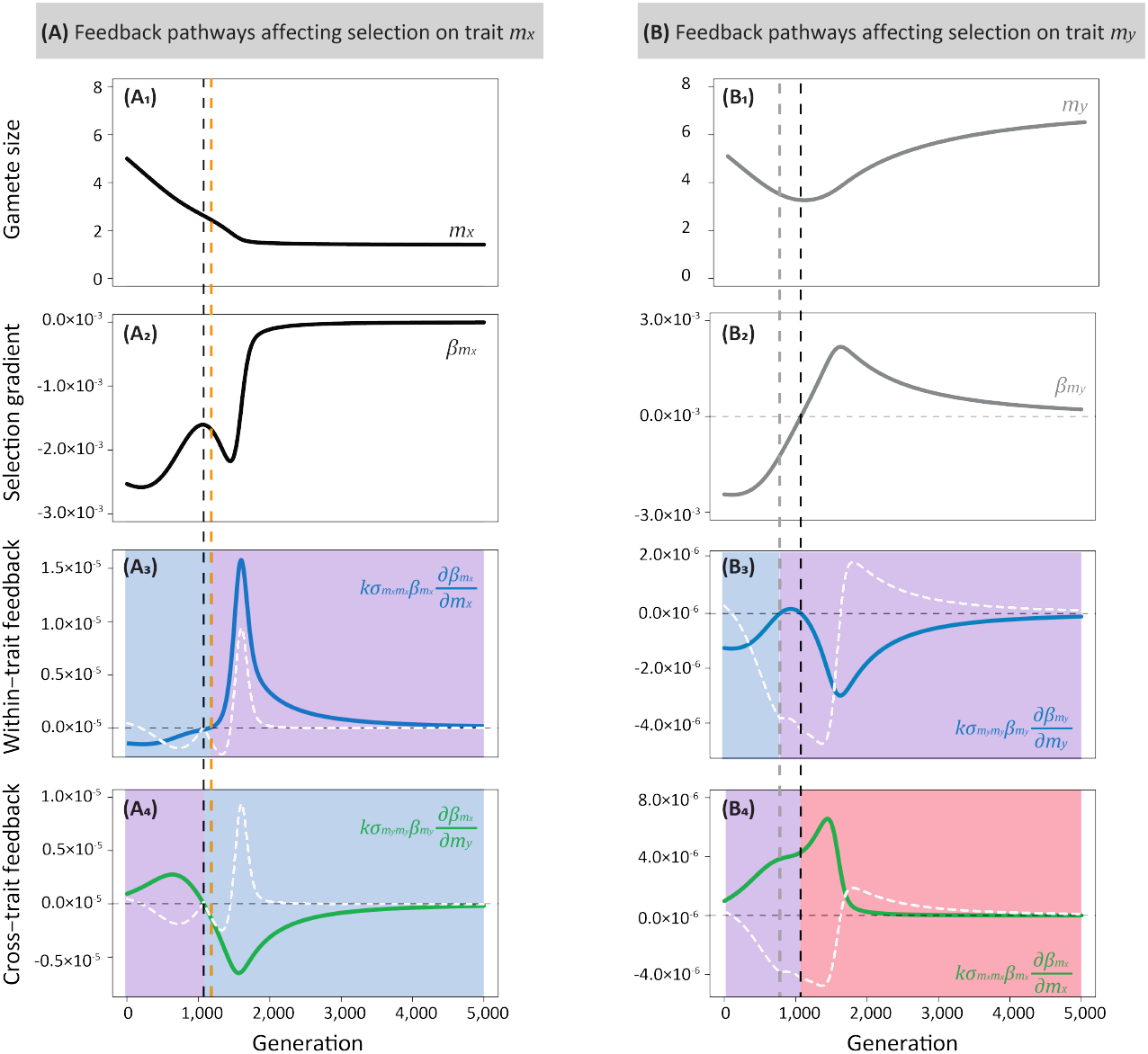
Application of feedback analysis to the evolution of anisogamy. Based on the model of Lehtonen nd Kokko (2011), gamete sizes of two mating types, *m*_*x*_ and *m*_*y*_, coevolve. **(A**_**1**_, **B**_**1**_) Evolutionary trajectories of *m*_*x*_ (black) and *m*_*y*_ (grey), following the canonical equation of adaptive dynamics. **(A**_**2**_, **B**_**2**_) Selection gradients 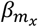 (black) and 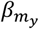 (grey) over evolutionary time. **(A**_**3**_, **B**_**3**_) Within-trait feedback (blue lines) on each selection gradient. **(A**_**4**_, **B**_**4**_) Cross-trait feedback (green lines) on each selection gradient. The white dashed lines represent the difference in absolute values between within- and cross-trait feedback components, where positive values indicate a greater magnitude of within-trait feedback. The vertical dashed lines (black, orange, and grey) indicate the time points at which the feedback components switch sign. The black vertical dashed lines also mark the time point when *m*_*y*_ transitions from a decreasing trend to an increasing one. The blue and red backgrounds indicate ‘negative reinforcing’ and ‘positive reinforcing’ feedback, respectively; and purple background shows all ‘inhibiting’ feedback. Parameter setting: *A*_*x*_ = *A*_*y*_ = 5, *M* = 1, *γ* = 1, *κ* = 1 and 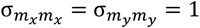. Gamete sizes *m*_*x*_ and *m*_*y*_ were initialized at 5 and 5.1, respectively. See detailed model description and other parameter values in Supplementary Materials. we approximated the evolution trajectories, selection gradients, and feedback components numerically using Wolfram Mathematica (see Mathematica code in Supplementary Materials).

Cross-trait feedback on 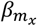 (i.e., changes in 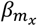 due to direct selection on *m*_*y*_) exhibits the opposite pattern to within-trait feedback. It begins as positive (left of the black dashed vertical line in Fig. 6A_4_) and switches to negative (right of the black dashed vertical line in Fig. 6A_4_). The switching point, marked by the black dashed vertical lines in Fig. 6, corresponds to a turning point for gametes with size *m*_*y*_, at which *m*_*y*_ shifts from a decreasing trend to an increasing one (Fig. 6B_1_) and the selection gradient 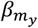 correspondingly switches from negative to positive (Fig. 6B_2_). Before this switching point, decreasing values of *m*_*y*_ act as a brake on selection for smaller *m*_*x*_ (i.e., negative inhibiting cross-trait feedback; purple background in Fig. 6A_4_), because the need to produce surviving zygotes prevents both gamete types from becoming too small simultaneously. After the switching point, increasing values of *m*_*y*_ accelerate selection for smaller *m*_*x*_ (i.e., negative reinforcing cross-trait feedback; blue background in Fig. 6A_4_). This is because larger values of *m*_*y*_ allow *m*_*x*_ to become even smaller without compromising the viability of zygotes. Within-trait and cross-trait feedback on 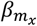 act in opposite directions over most of the evolutionary trajectory. By examining the difference between the absolute values of the within- and cross-trait feedback components (indicated by the white dashed lines in Figure 6A_3_, A_4_), we can see that inhibiting cross-trait feedback plays a more influential role in the initial evolution of *m*_*x*_. After *m*_*y*_ begins to increase, there is a brief period in which reinforcing cross-trait feedback dominates, causing selection for smaller *m*_*x*_ to increase. In the final phase, inhibiting within-trait feedback dominates as gamete sizes approach equilibrium.

Gametes of size *m*_*y*_, which are initially slightly larger than opposite-type gametes, experience an initial decrease in size followed by an increase (Fig. 6B_1_). Correspondingly, the selection gradient 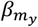 is initially negative, before becoming positive (Fig. 6B_2_). During the period of decreasing *m*_*y*_ (left of the black dashed vertical line in Fig. 6B), within-trait feedback follows a pattern very similar to that of *m*_*x*_. It initially exhibits reinforcing feedback (left of the grey dashed vertical line and in blue background in Fig. 6B_3_) and then transitions briefly to inhibiting feedback (between the grey and black dashed vertical lines and in purple background in Fig. 6B_3_). Cross-trait feedback is inhibiting throughout initial phase (Fig. 6B_4_). Once *m*_*y*_ starts to increase (right of the black dashed vertical line in Fig. 6B), within-trait feedback acts to inhibit selection for larger *m*_*y*_ (purple background in Fig. 6B_3_), whereas cross-trait feedback reinforces this selection (red background in Fig. 6B_4_). As with *m*_*x*_ above, inhibiting cross-trait feedback initially dominates the evolution of the selection gradient. Once *m*_*y*_ begins to increase, reinforcing cross-trait feedback dominates, causing increasing selection for larger *m*_*y*_. In the final phase, as the equilibrium is approached, inhibiting within-trait feedback exerts a greater influence, leading to a reduction in selection for larger *m*_*y*_.

Taken together, we gain a comprehensive understanding of the coevolutionary process involving the two gamete types. In the initial phase, both gamete types undergo a decrease in size. This reduction is strongly influenced by cross-trait feedback, which inhibits selection for smaller size in both gamete types, as the costs of reducing gamete size are greater if fertilisation partners produce smaller gametes. This aligns with previous arguments that sexual conflict is already present in the earliest stages of the evolution towards anisogamy (Lehtonen et al. 2012). The initially smaller gametes decrease more rapidly, reaching a size that halts further reduction in the opposite type; otherwise, the zygote is unlikely to survive. As the initially smaller gametes continue to decrease in size while the opposite type begins to increase in size, they reinforce the strength of selection on each other via cross-trait feedback, leading to further asymmetry in their sizes. At the end of the coevolutionary process, within-trait feedback becomes more dominant, reducing the strength of selection and causing each type of gamete to evolve slowly toward its equilibrium state.

### Example 3: Fisherian runaway process of sexual selection

Our final example, based on the Fisherian runaway process of sexual selection, illustrates the application of feedback analysis in quantitative genetic models under the weak selection limit. The foundational model for the Fisher process was developed by Lande (1981) (but see Henshaw & Jones, 2020). Lande’s model can be easily solved under the weak selection limit, yielding the same mathematical predictions. In this model, populations are characterized by two quantitative traits: a male ornament *s* and a female preference *p*. Both traits are inherited via autosomal transmission and exhibit sex-specific expression. Assuming weak selection, the distributions of male ornament and female preference can take any form with means 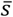 and 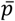, provided that both traits are tightly concentrated around their respective means. The model assumes no direct benefits or costs for female choosiness and thus, the selection gradient on female preference is zero. However, indirect selection on preference occurs due to its additive genetic correlation with male ornamentation. Therefore, the changes in 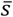 and 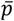 from the current to the next generation can be described as 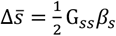 and 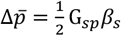, respectively (cf. equation (13)). The factor 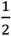 accounts for sex-specific expression, *G*_*ss*_ represents the additive genetic variance for male ornament size, *G*_*sp*_ is the additive genetic covariance between male ornament size and female preference strength, and *β*_*s*_ is the selection gradient for the male ornament. In the model of Lande (1981), the fitness of a male with ornament size *s* in a population with mean preference 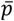 is given by 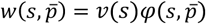. Here, the viability function 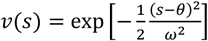 determines the survival probability of a male with ornament size *s* (where 1/*ω* reflects the strength of viability selection against exaggerated ornaments and *θ* measures the optimal ornament size under viability selection). The ‘psychophysical’ preference function 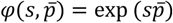 determines the mating success of a male with ornament size *s* in a population with the mean female preference strength 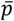. Females with *p* > 0 prefer males with more extreme ornamentation. The selection gradient for male ornament is therefore given by as 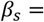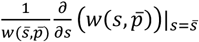. Setting the selection gradient equal to zero yields a line of equilibria for female preference and male ornament: 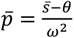, with the slope given by 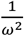.

By applying equation (15), we can analyse how changes in male ornamentation and female preference influence selection on 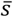. There are two feedback components at play here: within-trait feedback (given by 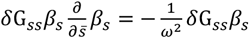), and cross-trait feedback due to indirect selection on female preference (given by 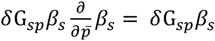). For a better understanding, we here focus on the case where the male ornament is initialized with a value smaller than the optimal size (*s* < *θ*), and thus, there is selection favouring larger male ornament (*β*_*s*_ > 0). Accordingly, the within-trait feedback 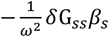 becomes negative, suggesting that as male ornament size increases, selection for further enlargement diminishes. This self-inhibiting phenomenon arises due to male ornamentation approaching its optimal size for a fixed preference strength. On the other hand, the cross-trait feedback *δ*G_*sp*_*β*_*s*_ is positive, indicating a positive reinforcing effect. This means that as female preference increases, selection for even larger male ornaments is enhanced.

By comparing the magnitudes of each feedback component 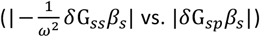, we can understand how they interact with the selection gradient (note that we focus on the case of *β*_*s*_ > 0), thereby influencing the evolution of male ornamentation and female preference. When the within-trait feedback is larger in magnitude 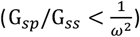, selection for larger male ornamentation is predominantly inhibiting until it reaches the equilibrium. This inhibiting effect extends to the genetically correlated female preference, resulting in a similar evolutionary trajectory. Therefore, in this case the evolution will proceed towards the line of equilibria with slope 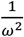. In contrast, when the cross-trait feedback has is larger in magnitude 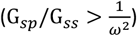, increases in female preference further amplify selection for even larger ornament in males, resulting in a never-ending runaway process. This process accelerates rapidly away from any equilibrium point, with traits evolving at an ever-increasing speed. Our framework therefore provides a new lens through which to view Lande’s (1981) original analysis.

## Discussion

In this study, we provide a general framework for quantifying how changes in coevolving traits, caused by selection, feed back to shape the selection environment. This framework can be applied both to adaptive-dynamic models and to quantitative-genetic models that assume weak selection. Our framework can be used to provide formal support or refutation of verbal arguments about exactly why certain traits coevolve in the manner they do.

The application of our framework to the evolution of anisogamy (Example 2) illustrates its power for interrogating verbal models of coevolution. In the model we analyze, small initial asymmetries in gamete size can amplify over evolutionary time, leading to divergence into small gametes (sperm) and large gametes (eggs). Trivers (1972) famously argued that any initial differences in parental investment between the sexes are maintained or magnified over time a via self-reinforcing process. Trivers’ argument was as follows: Because females initially invest more than males in zygotes, they have more to lose if they abandon their offspring and so should be incentivized to provide further parental investment (e.g., larger eggs or post-zygotic care). In contrast, males, who invest less in gametes and consequently become mate-limited, should allocate more resources to competitive traits to enhance their mating success, rather than increasing parental investment. This argument has been criticized for various reasons (e.g., it commits the ‘Concorde fallacy’; see Dawkins & Carlisle, 1976; Kokko & Jennions, 2008). Some authors argue that anisogamy cannot be the fundamental cause of sex-role divergence (Gowaty & Hubbell, 2005; Schärer et al., 2012), while others have suggested that anisogamy can drive sex-role divergence when other factors are taken into account (e.g., Queller, 1997; Fromhage & Jennions, 2016).

Our feedback analysis of the evolution of anisogamy suggests a new take on Trivers’ argument: In our analysis, sex differences in gamete investment are not maintained by a self-reinforcing process (‘I should invest more because I already invested more’), but rather by a cross-trait reinforcing process (‘I should invest more because my partner invested less’ and ‘I should invest less because my partner invested more’). Importantly, reinforcement here occurs over evolutionary time, rather than within a generation. As depicted in Fig. 6, if one gamete type is larger than the other, the selection environment for each gamete type is strongly shaped by cross-trait reinforcing feedback. The decreasing size of the smaller gamete creates a selection environment that favours even larger size in the larger gamete. Similarly, the increasing size of the larger gamete selects for further reductions in the size of the smaller gamete. In contrast to Trivers’ argument, changes in the size of one gamete type often lead to *weaker* selection for further changes in the same direction (within-trait inhibiting feedback). An analogous argument could potentially be developed to explain the divergence of post-zygotic parental investment (cf. McNamara et al. 2003), the main target of Trivers’ original argument, although careful consideration would be needed to account for the differences in the selection pressures acting on gamete size and subsequent parental investment.

To apply our framework in the context of adaptive-dynamic or quantitative-genetic models, a key assumption must be satisfied: The individual fitness surface must be approximately linear over the range of trait variation. This restriction is required because we approximate the fitness surface using a first-order Taylor polynomial (i.e., a linear approximation) when estimating how selection gradients change due to trait evolution. Typically, this assumption is framed as ‘weak selection’, since it is met for any smooth fitness surface when trait values are tightly distributed around the mean trait value (Wild & Traulsen, 2007; McElreath & Boyd, 2008). However, this assumption is also consistent with strong selection, as long as the fitness surface is nearly linear over the range of trait variation. Under strong and non-linear selection, however, our feedback framework should be viewed only as an approximation. This is because, in such cases, the feedback components (i.e., within- and cross-trait feedback as demonstrated in our study) do not interact additively, posing challenges in disentangling the contribution of each feedback component to changes in selection environment. Given the key assumptions of our framework, our feedback analysis could feasibly be extended to population genetic models, particularly in scenarios involving weak selection (Barton & Turelli, 1991; Kirkpatrick et al., 2002; van Doorn & Kirkpatrick, 2010). However, the genetic details, including factors such as dominance and epistasis, along with interactions between multiple genetic loci, can lead to more complex coevolutionary dynamics. Therefore, it may be necessary to consider these factors when extending our feedback analysis in this context.

Lastly, our framework assumes that mean trait values alone are sufficient to determine evolutionary dynamics. This simplification is insufficient to capture some important classes of evolutionary dynamics, however. For example, in some adaptive-dynamic models, the dynamics lead to evolutionary branching (Geritz et al., 1998). Branching corresponds to an increase in phenotypic variance that results in polymorphic populations with a bimodal distribution of traits (Geritz et al., 1998; Dercole & Rinaldi, 2008). As a consequence, our approach as developed here cannot provide insights once traits converge to a branching point. Further, both the **M**-matrix and the **G**-matrix may evolve, rather than being fixed as we assume here (Blows, 2007; Jones et al., 2007; Arnold et al., 2008). This in turn may feed back to influence the strength and direction of selection. While the framework developed here assumes that selection is sensitive to trait means only, future work could extend this framework to consider how changes in trait distributions more broadly shape the selection environment.

## Data and code availability

The Mathematica scripts for the examples presented in this study will be available in the Supplementary Materials.

## Author contributions

X.L. and J.M.H. came up with the original idea; X.L. and J.M.H. developed the methods; All three authors contributed by discussing theory, methods and concepts. X.L. wrote the manuscript; All three authors revised the manuscript.

## Funding

This work was funded by the Bundesministerium für Bildung und Forschung, Germany, and the Deutsche Forschungsgemeinschaft (DFG, German Research Foundation) under project number 456626331. J.L. was funded by an Academy of Finland grant (340130).

## Conflict of interest statement

No competing interest is declared.

## Notes

### Competing Interest Statement

The authors have declared no competing interest.

## References

Andersson, M. (1994). Sexual Selection. Princeton University Press, Princeton.

Arnold, S. J., Bürger, R., Hohenlohe, P. A., Ajie, B. C., & Jones, A. G. (2008). Understanding the evolution and stability of the G-matrix. Evolution, 62(10), 2451–2461.

Baldauf, S. A., Engqvist, L., & Weissing, F. J. (2014). Diversifying evolution of competitiveness. Nature Communications, 5(1), 5233.

Bank, C. (2022). Epistasis and adaptation on fitness landscapes. Annual Review of Ecology, Evolution, and Systematics, 53(1), 457–479.

Barton, N. H., & Turelli, M. (1991). Natural and sexual selection on many loci. Genetics, 127(1), 229– 255.

Blows, M. W. (2007). A tale of two matrices: multivariate approaches in evolutionary biology. Journal of evolutionary biology, 20(1), 1–8.

Champagnat, N., Ferriere, R., & Ben Arous, G. (2001). The canonical equation of adaptive dynamics: a mathematical view. Selection, 2, 71–81.

Dashtbali, M., Long, X., & Henshaw, J. M. (2024). The evolution of honest and dishonest signals of fighting ability. Evolution Letters, qrae008.

Dawkins, R., & Carlisle, T. R. (1976). Parental investment, mate desertion and a fallacy. Nature, 262(5564), 131–133.

Geritz, S. A., Kisdi, E., Meszé NA, G., & Metz, J. A. (1998). Evolutionarily singular strategies and the adaptive growth and branching of the evolutionary tree. Evolutionary ecology, 12, 35–57.

Dieckmann, U., & Law, R. (1996). The dynamical theory of coevolution: a derivation from stochastic ecological processes. Journal of mathematical biology, 34, 579–612.

Fawcett, T. W., Kuijper, B., Weissing, F. J., & Pen, I. (2011). Sex-ratio control erodes sexual selection, revealing evolutionary feedback from adaptive plasticity. Proceedings of the National Academy of Sciences, 108(38), 15925–15930.

Fromhage, L., & Jennions, M. D. (2016). Coevolution of parental investment and sexually selected traits drives sex-role divergence. Nature communications, 7(1), 12517.

Dercole, F., & Rinaldi, S. (2008). Analysis of evolutionary processes: the adaptive dynamics approach and its applications: the adaptive dynamics approach and its applications. Princeton University Press, Princeton.

Giraldeau, L. A., & Caraco, T. (2000). Social foraging theory (Vol. 73). Princeton University Press, Princeton.

Gowaty, P. A., & Hubbell, S. P. (2005). Chance, time allocation, and the evolution of adaptively flexible sex role behavior. Integrative and Comparative Biology, 45(5), 931–944.

Harten, L., Matalon, Y., Galli, N., Navon, H., Dor, R., & Yovel, Y. (2018). Persistent producer-scrounger relationships in bats. Science Advances, 4(2), e1603293.

Henshaw, J. M., Fromhage, L., & Jones, A. G. (2019). Sex roles and the evolution of parental care specialization. Proceedings of the Royal Society B, 286(1909), 20191312.

Henshaw, J. M., & Jones, A. G. (2020). Fisher’s lost model of runaway sexual selection. Evolution, 74(2), 487–494.

Henshaw, J. M., Fromhage, L., & Jones, A. G. (2022). The evolution of mating preferences for genetic attractiveness and quality in the presence of sensory bias. Proceedings of the National Academy of Sciences, 119(33), e2206262119.

Hochberg, M. E., Rankin, D. J., & Taborsky, M. (2008). The coevolution of cooperation and dispersal in social groups and its implications for the emergence of multicellularity. BMC Evolutionary Biology, 8, 1–14.

Iwasa, Y., Pomiankowski, A., & Nee, S. (1991). The evolution of costly mate preferences II. The “handicap” principle. Evolution, 45(6), 1431–1442.

Jones, A. G., Arnold, S. J., & Bürger, R. (2007). The mutation matrix and the evolution of evolvability. Evolution, 61(4), 727–745.

Kirkpatrick, M. (1982). Sexual selection and the evolution of female choice. Evolution, 1–12.

Kirkpatrick, M., Johnson, T., & Barton, N. (2002). General models of multilocus evolution. Genetics, 161(4), 1727–1750.

Kuijper, B., Pen, I., & Weissing, F. J. (2012). A guide to sexual selection theory. Annual Review of Ecology, Evolution, and Systematics, 43(1), 287–311.

Lande, R. (1981). Models of speciation by sexual selection on polygenic traits. Proceedings of the National Academy of Sciences, 78(6), 3721–3725.

Lande, R., & Arnold, S. J. (1983). The measurement of selection on correlated characters. Evolution, 1210–1226.

Lehtonen, J., & Kokko, H. (2011). Two roads to two sexes: unifying gamete competition and gamete limitation in a single model of anisogamy evolution. Behavioral ecology and sociobiology, 65, 445–459.

Lehtonen, J., Jennions, M. D., & Kokko, H. (2012). The many costs of sex. Trends in ecology & evolution, 27(3), 172–178.

Leimar, O. (2009). Multidimensional convergence stability. Evolutionary Ecology Research, 11(2), 191– 208.

Long, X., & Weissing, F. J. (2023). Transient polymorphisms in parental care strategies drive divergence of sex roles. Nature Communications, 14(1), 6805.

Long, X., Komdeur, J., Székely, T., & Weissing, F. J. (2024). A life-history perspective on the evolutionary interplay of sex ratios and parental sex roles. American Naturalist. In press.

Mathot, K. J., & Giraldeau, L. A. (2008). Increasing vulnerability to predation increases preference for the scrounger foraging tactic. Behavioral Ecology, 19(1), 131–138.

McElreath, R., & Boyd, R. (2008). Mathematical models of social evolution: A guide for the perplexed. University of Chicago Press.

McGill, B. J., & Brown, J. S. (2007). Evolutionary game theory and adaptive dynamics of continuous traits. Annual Review of Ecology, Evolution, and Systematics, 38(1), 403–435.

McNamara, J. M., & Leimar, O. (2020). Game theory in biology: concepts and frontiers. Chapter 6. Oxford University Press, USA.

Mead, L. S., & Arnold, S. J. (2004). Quantitative genetic models of sexual selection. Trends in ecology & evolution, 19(5), 264–271.

Mullon, C., Keller, L., & Lehmann, L. (2018). Social polymorphism is favoured by the co-evolution of dispersal with social behaviour. Nature ecology & evolution, 2(1), 132–140.

Nagylaki, T. (1992). Introduction to theoretical population genetics (Vol. 21). Springer Science & Business Media, New York.

Pen, I., & Weissing, F. J. (2001). Sexual selection and the sex ratio: an ESS analysis. Selection, 1(1-3), 111–122.

Pigliucci, M. (2006). Genetic variance–covariance matrices: a critique of the evolutionary quantitative genetics research program. Biology and Philosophy, 21, 1–23.

Powers, S. T., Penn, A. S., & Watson, R. A. (2011). The concurrent evolution of cooperation and the population structures that support it. Evolution, 65(6), 1527–1543.

Queller, D. C. (1997). Why do females care more than males? Proceedings of the Royal Society of London. Series B: Biological Sciences, 264(1388), 1555–1557.

Ranta, E. S. A., Peuhkuri, N., Hirvonen, H., & Barnard, C. J. (1998). Producers, scroungers and the price of a free meal. Animal Behaviour, 55(3), 737–744.

Revathi Venkateswaran, V., Roth, O., & Gokhale, C. S. (2021). Consequences of combining sex-specific traits. Evolution, 75(6), 1274–1287.

Roff, D. A. (1997). Evolutionary quantitative genetics. Springer Science & Business Media, New York.

Schärer, L., Rowe, L., & Arnqvist, G. (2012). Anisogamy, chance and the evolution of sex roles. Trends in Ecology & Evolution, 27(5), 260–264.

Smith, C. C., & Fretwell, S. D. (1974). The optimal balance between size and number of offspring. The American Naturalist, 108(962), 499–506.

Svensson, E. I., Arnold, S. J., Bürger, R., Csilléry, K., Draghi, J., Henshaw, J. M., … & Runemark, A. (2021). Correlational selection in the age of genomics. Nature ecology & evolution, 5(5), 562–573.

Taylor, P. D. (1996). The selection differential in quantitative genetics and ESS models. Evolution, 2106–2110.

Trivers, R. L. (1972). Parental investment and sexual selection. In Sexual Selection and the Descent of Man 1871–1971 (eds Houck, L. D. & Drickamer, L. C.. University of Chicago Press.

van Doorn, G. S., & Kirkpatrick, M. (2010). Transitions between male and female heterogamety caused by sex-antagonistic selection. Genetics, 186(2), 629–645.

Waffender, A., & Henshaw, J. M. (2023). Long-term persistence of exaggerated ornaments under Fisherian runaway despite costly mate search. Journal of Evolutionary Biology, 36(1), 45–56.

Wild, G., & Traulsen, A. (2007). The different limits of weak selection and the evolutionary dynamics of finite populations. Journal of Theoretical Biology, 247(2), 382–390.

