## Supplementary Materials for "Quantifying feedback among traits in coevolutionary models"

**Example 2: The evolution of anisogamy**

In the model of Lehtonen and Kokko (2011), they assume that the initially resident population is isogamous, consisting of two different mating types, denoted as $x$ and $y$. These two mating types produce gametes with size, $m_x$ and $m_y$, respectively. While both gametes share similar sizes, disassortative fusion of them necessitated to produce zygotes. In a local mating group, there are fixed number of $A_x$ and $A_y$ adult individuals with the mating type $x$ and $y$, respectively. According to their model, an increase in the values of $A_x$ and $A_y$ corresponds to greater gamete competition. Additionally, their model assumes no sex ratio bias, meaning that $A_x$ is always equal to $A_y$. For each adult individual, there is a fixed amount of resource $M$ for producing gametes per time unit. Therefore, each mating type can produce a total number of $A_x \frac{M}{m_x}$and $A_y \frac{M}{m_y}$ gametes per time unit in a local group. Moreover, gametes produced by the two mating types experience instantaneous mortality rates of $\mu(m_x) = \varphi \exp{(\frac{\rho}{m_x})^k}$ and $\mu(m_y) = \varphi \exp{(\frac{\rho}{m_y})^k}$, respectively; where $\varphi$ determines the minimum mortality rate, $\rho$ approximates the minimum gamete size, and $k$ influences how quickly the mortality rate increases as gamete size approaches the minimum threshold($\approx \rho$). The surviving gametes have chances of being fertilized. For each mating type, the overall fertilization rate is given by $\gamma N_x N_y$, where $\gamma$ is a fixed parameter determining the encounter rate of different gametes, and $N_x$ and $N_y$ are the numbers of gametes available for fertilization per time unit from mating types type $x$ and $y$, respectively. Taken together, the dynamics of $N_x$and $N_y$ are influenced by the reproduction of adult individuals, instantaneous mortality and fertilization rates:

$$\frac{dN_x}{dt} = A_x \frac{M}{m_x} - \mu(m_x)N_x - \gamma N_x N_y \quad (S1a)$$

$$\frac{dN_y}{dt} = A_y \frac{M}{m_y} - \mu(m_y)N_y - \gamma N_x N_y. \quad (S1b)$$

Setting equations $(S1a)$ and $(S1b)$ equal to zero allows us to determine the equilibrium number of each gamete available for fertilization. Now, the fitness for each mating type, defined as the total number of surviving zygotes, can be derived using the equilibrium values of $N_x$ and $N_y$. For a focal resident individual of mating type $x$ producing gametes with size $m_x$, its fitness in the resident population is given by $w(m_x, m_y) = \gamma N_x N_y S(m_x, m_y)$, where $S(m_x, m_y) = \exp\left(-\frac{\alpha}{m_x+m_y}\right)$ represents the zygote survival probability, with α scaling the relationship between zygote size and its survival. Accordingly, the larger the zygotes, the greater the probability of their survival. Moreover, the fitness of a focal resident individual of mating type $y$ is equal to that of mating type $x$ according to Fisher condition.

To know how gamete sizes, $m_x$ and $m_y$, evolve, one needs to consider a single mutant individual with either mating type, occurring in the resident population. For example, a mutant individual of mating type $x$ can arise. This mutant produces gametes of size $\widehat{m}_x$ and contributes a number of $\widehat{N}_x$ gametes available for fertilization in the population. Due to the interactions between the mutant and resident population, the dynamics of $N_x$, $N_y$ and $\widehat{N}_x$ can be described as:

$$\frac{dN_x}{dt} = (A_x - 1)\frac{M}{m_x} - \mu(m_x)N_x - \gamma N_x N_y \quad (S2a)$$

$$\frac{dN_y}{dt} = A_y\frac{M}{m_y} - \mu(m_y)N_y - \gamma(N_x + \widehat{N}_x)N_y \quad (S2b)$$

$$\frac{d\widehat{N}_x}{dt} = \frac{M}{\widehat{m}_x} - \mu(\widehat{m}_x)\widehat{N}_x - \gamma \widehat{N}_x N_y. \qquad (S2c)$$

Again, the equilibrium values of $N_x$, $N_y$ and $\widehat{N}_x$ can be obtained by equating equations $(S2a)$, $(S2b)$ and $(S2c)$ to zero. As a result, the invasion fitness of the mutant which produces gametes of size $\widehat{m}_x$ is given by

$$w(\widehat{m}_x, m_x, m_y) = \gamma \widehat{N}_x N_y S(\widehat{m}_x, m_y) \quad (S3)$$

When a mutant individual of mating type $y$ (producing gametes with size $\widehat{m}_y$) emerges, its invasion fitness $w(\widehat{m}_y, m_x, m_y)$ can be determined by substituting all $x$- and $y$ - indices in equations $(S2)$ and $(S3)$. Based on the invasion fitness, we can derive the selection gradients for each mating type:

$$\beta_{m_x} = \frac{1}{w(m_x, m_y)}\frac{\partial}{\partial \widehat{m}_x} w(\widehat{m}_x, m_x, m_y)|_{\widehat{m}_x = m_x} \quad (S4a)$$

$$\beta_{m_y} = \frac{1}{w(m_x, m_y)} \frac{\partial}{\partial \widehat{m}_y} w(\widehat{m}_y, m_x, m_y)|_{\widehat{m}_y = m_y}. \quad (S4b)$$

If you are interested, please refer to the original paper by Lehtonen and Kokko (2011) for detailed calculus and solutions of the selection gradient.

In Figure 6, we use the parameter setting: $A_x = A_y = 5$, $M = 1$, $\varphi = 1$, $\rho = 1$ , $k = 5$, $\gamma = 1$, $\alpha = 10$, $\kappa\sigma_{m_x m_x} = \kappa\sigma_{m_y m_y} = 1$. As demonstrated in Lehtonen and Kokko (2011), with this parameter setting, an intermediate level of gamete competition is assumed ($A_x = A_y > 1$). An initial small asymmetry in gamete sizes is sufficient to initiate the evolution of anisogamy when $\alpha > 4\rho$.
